# Structural variation detection with read pair information — An improved null-hypothesis reduces bias

**DOI:** 10.1101/036707

**Authors:** Kristoffer Sahlin, Mattias Frånberg, Lars Arvestad

## Abstract

Reads from paired-end and mate-pair libraries are often utilized to find structural variation in genomes, and one common approach is to use their fragment length for detection. After aligning read-pairs to the reference, read-pair distances are analyzed for statistically significant deviations. However, previously proposed methods are based on a simplified model of observed fragment lengths that does not agree with data. We show how this model limits statistical analysis of identifying variants and propose a new model, by adapting a model we have previously introduced for contig scaffolding, which agrees with data. From this model we derive an improved improved null hypothesis that, when applied in the variant caller CLEVER, reduces the number of false positives and corrects a bias that contributes to more deletion calls than insertion calls. A reference implementation is freely available at https://github.com/ksahlin/GetDistr.

## 1 Introduction

Genomic structural variation, for example insertion and deletion of DNA, are common in the human population and have been linked to various diseases and conditions. The basic question scientists and clinicians want to answer is: given a DNA sample from a donor and a suitable reference genome, what structural variants does the donor have in comparison to the reference? Methods for identifying structural variants are continuously worked on, in terms of both experimental protocols and bioinformatic analysis. Short-read technologies are, despite their weaknesses, the primary data source because of the superior throughput/cost ratio. It is today important to improve accuracy of predictions and in particular to reduce the false-positive rate while retaining sensitivity. To that end, we have worked on improving the statistical analysis of paired reads, using paired-end (PE) or mate-pair (MP) libraries, for evaluating the significance of a detected insertion or deletion.

While aligned reads are important for identifying short variants and substitutions, larger variants and variants in repetitive regions where alignment is difficult are easier detected by paired reads spanning over the region. In PE and MP protocols, reads are from the ends of DNA *fragments* from the donor. PE libraries have short-range fragment lengths (up to 100s bp), MP libraries are long range (1000s bp), and they each have their own strengths and limitations. PE libraries often has superior coverage and narrow fragment length distribution while long range MP libraries can span larger insertions and, at similar read coverage, provide higher span coverage (the number of MP pairs separated by a random position) than PE libraries, which in theory can make up for the increased variation in individual fragment lengths by increasing statistical power from more observations.

### 1.1 Previous work

Numerous structural variation algorithms using read pairs, and their fragment length, to detect variants have been proposed. Many tools use only *discordant* read pairs for downstream calling of variants, i.e., read pairs that align at a distance smaller than 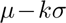 or larger than 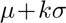 base pairs from each other, where *μ* and *σ* are the mean and standard deviation of the fragment length distribution and 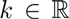 [2,5,11,12,20]. This restriction may reduce the computational demand, but it sacrifices sensitivity [17] by removing observations.

There are also tools with a statistical model/approach that utilizes all read pairs. CLEVER [17] finds insertions and deletions based on statistically significant deviation of the mean fragment length of all reads^5^ over a position from μ. This method finds more and smaller variants compared to methods that use only discordant reads [17]. [9] models the number of discordant and concordant read pairs (classified by a cutoff) over a region as following a binomial distribution and finds inversions and deletions based on statistically significant accumulation of discordant read pairs over regions. However, any binary classification cutoff causes loss of information [8], thus statistical power, as they do not consider how much above or below the cutoff a fragment length is^6^. Another approach is non-parametric testing of the distribution over a region, e.g., using the Kolmogorov-Smirnov test [15], but as [17] noted, this is computationally expensive. [10] presented a model to find the most likely common deletion length from several donor genomes with different fragment length distributions by maximizing the likelihood of observed fragment lengths given a deletion size and each of the distributions.

These methods however assumes that the probability of a fragment length being observed over a position/region follows the probability distribution of the full library fragment length distribution, which is not true [22]. Longer fragment lengths span more positions than shorter fragment lengths, so over any position in the genome there will be a bias towards read-pairs further apart than *μ*. This observation bias of fragment sizes has been investigated earlier in an assembly context, estimating the gap size between contigs [22,21,18]. The approaches given in [21,18] are more general by using the exact (empirical) distribution over the fragment length, which also makes them computationally demanding. GapEst [22] assumes a normal fragment length distribution and derives an analytic expression for the likelihood of a gap size that scales very well, which opens up for other applications where this type of problem needs to be calculated for a large number of instances, *e.g.*, structural variation detection. There is no previous work known to the authors on incorporating this model, or a similar one, to structural variation and investigating how it affects the balance between detecting deletions and insertions.

### 1.2 Contribution

We use the statistical model given in [22] and present it in the context of structural variation detection. The model provides a probability distribution for the fragment sizes we observe over a position (*e.g.*, a potential *breakpoint*) or region. Given this distribution we derive a null-hypothesis distribution to detect variants. We show that the corrected null-hypothesis agrees with both simulated and biological data, while a commonly used null-hypothesis does not. We implement the null-hypothesis in the state-of-the-art fragment-length based variant caller CLEVER [17]. Although CLEVER uses constraints and assumptions that do not agree with our model, we show that the detection of insertions and deletions becomes more balanced and that the number of false positive calls decreases. This is a promising first result as we could only apply a part of our theory in CLEVER without a significant restructuring of the code. We also believe that this work is a step towards creating a statistical rigorous approach for read pair fragment lengths where we can detect indels to a much higher resolution than cutoff based ones.

## 2 Methods

We will review a model used to determine contig distances in scaffolding [22] and use it in the context of structural variation detection. Notation and assumptions are presented in section 2.1. In section 2.2 we present the probability distribution in a structural variation detection context. Section 2.4 discuss a commonly used null-hypothesis used for detecting variants with fragment length and derives an improved null-hypothesis using our model. Some text is deferred to an Appendix.

### 2.1 Notation and assumption

We refer to our model as the *Observed Fragment Length* (OFL) model. This model carries no new concepts and makes the same assumptions as the Lander Waterman model [14], but adds a variable and some constants. We only state it here for convenience of referencing to a model when we derive probabilities and a null-hypothesis. Read pairs are sampled independently and uniformly from the donor genome. Let *G* denote the length of the reference genome. Alignment of read pairs to the reference genome yields our observations: distance *o* between reads in a read pair, read length r, and number of allowed “inner”^7^ softclipped bases *s* [16] in an alignment, see Figure 1a. Read-pair distances *x* come from a library fragment length distribution *f*(*x*) (either given or estimated from alignments). We denote the mean and standard deviation of this distribution as *μ* and *σ*. Finally, a parameter *δ* models the number of missing or added base pairs in the reference, compared to the donor sequence. That is, if the donor sequence contains an insertion, *δ* is negative and we say that the donor sequence has *δ* added bases. Similarly, if the donor sequence contains a deletion, *δ* is positive and we say that the donor sequence has *δ* deleted bases. For a given read-pair with fragment length *x*, 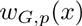 denote the probability that it spans over position *p* on a genome of size *G*. As we do not model that any two positions have different probabilities to be spanned over (reads are drawn uniformly), *w* will not depend on *p* and we omit it and refer to 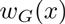 from now on.

**Fig. 1:**
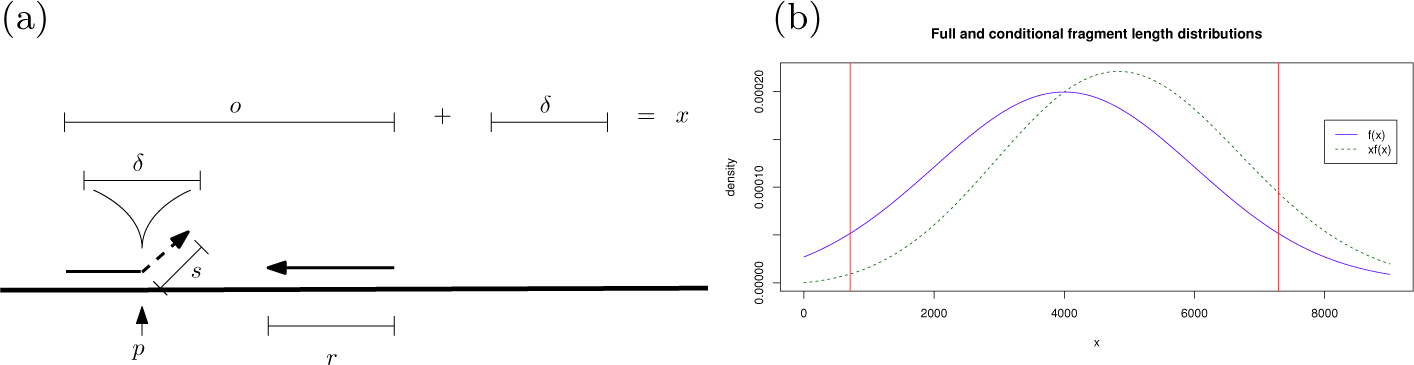
(a) Constants and variables in the OFL model. The figure illustrates the scenario of an insertion in the donor genome of length *δ* at position *p*. Two reads (marked by arrows) of length *r* are at distance *o* from each other, with the left read partially aligned leaving *s* positions unaligned (softclipped). (b) Illustration of a full fragment size distribution *f*(*x*) from *N*(4000, 2000) (blue line), from which *H*_0_ is derived. The green dashed line shows the observed fragment distribution over any variant free position for fragments coming from *f*(*x*) for the simplified case when 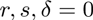 (i.e., this is exactly the function 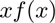. Red lines indicate the 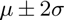 quantiles of *f*(*x*). It is less likely to observe a smaller fragment size over any given position in the genome (see density of green distribution at red lines), as opposed to identical significance under *f*(*x*).

### 2.2 Probability function over observed fragment lengths

The distribution and probabilities derived in this section closely matches those given in [22] with the minor addition of the constant *s*. We restate the expressions in a structural variation context for clarity.

*No variant* First, we assume that donor and reference are identical, therefore 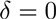 at any position. Given the OFL model, the probability that we observe a read pair with fragment size *x* over a position *p* on a genome of length *G* is

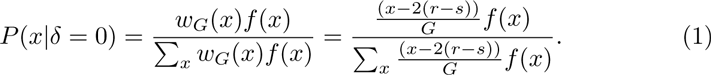

Here *f*(*x*) is the probability to draw a fragment of length *x* from the full library and 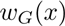 the probability that it spans over position *p*. The denominator is a normalization constant to make *P* a probability. It is assumed that 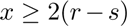 For example, if the read length is 100 and maximum allowed softclipped bases of an aligned read is 30 a read pair with fragment length 300 will have 300 − 2(100 − 70) = 160 possible placements where it spans position *p*. For simplicity, we omit the special case when *p* is near the end of a chromosome.

*Modeling variant at a position* Let *δ* be the unknown variant size. In this case we cannot observe the true fragment length *x* of read pairs. What we observe is instead 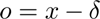 (see Figure 1a). A modification of *w*(*x*) is needed as fragment sizes is now required to span *δ* base pairs and have sufficiently many base pairs on each side to be mapped 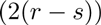. We have

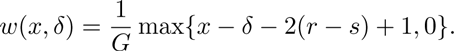

The 0 in the max function keeps the function weight to 0 in case we have no possible placing of a paired read over a variant. We can simplify this function to be expressed in *o*, as 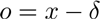, and write 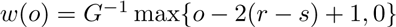.

We see that the function *w* is constant for any given observation and can therefore be interpreted as a “weight” function, hence the notation *w*.

### 2.3 Probability of variant size *δ*

We can express the probability of *δ* given observations as 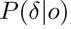. Lacking prior information about *δ*, we model it with the uniform distribution^8^. Using Bayes theorem, we get

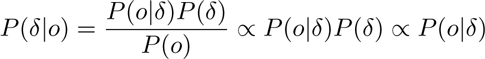

where 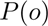 and 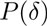 are constant by the assumption of a uniform distribution. We now have

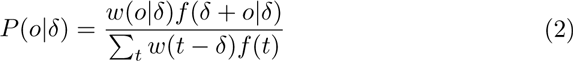

where the denominator is the sum of all possible fragment sizes that can be observed given *δ* and *f*. We can now find the most likely *δ* using maximum likelihood estimation (MLE) over (2). The time complexity for the MLE is 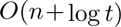^9^ if 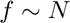 (with *t* continuous), where *n* is the number of observations [22]. Note that we implicitly get 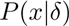 since 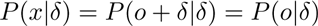.

### 2.4 Null-hypothesis and statistical testing

Let 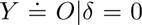, that is, the random variable over observed fragment lengths given 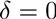. Let 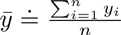 be the sample mean of observed fragment lengths. Considering 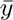 a random variable over experiments, it is commonly assumed that 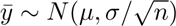 *i.e.*, the distribution of the sample mean of *f*(*x*) under the central limit theorem (CLT), and this distribution is used as null-hypothesis [5,17]. We call this null-hypothesis *H*_0_. Furthermore, the variant size *δ* is estimated from observed fragment lengths *o* as 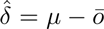 [5,17,9,12,20,10]. At first glance this formula seems reasonable since we take the expected fragment size and subtract the mean of the observations, but it has strong limitations. One is that 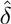in this case has an upper bound of 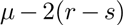 since 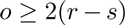. This equation implies that we can never span over a sequence longer than 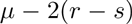. We use Equation 2 to derive the correct mean and standard deviation of *Y* given the OFL model, denoted 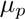 and 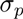 respectively. The derivation of 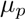 is similar to derivation of observed fragment size linking two contigs given in [22], and the derivation of 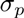 is a special case of the derivation of the variance of observed fragment size linking two contigs given in [23]. See proof in Appendix 5.5.

**Theorem 1**. *Given the OFL model*, 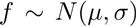, and 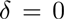, we have 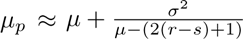 and 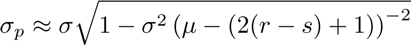.

The null-hypothesis is that there is no variant, thus 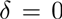. Under CLT, as *n* increases, we therefore have 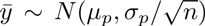. Notice that we can calculate 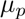 and 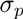 without the assumption 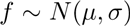 by using an empirical estimate of *f*(*x*) from aligned read pairs. Nevertheless, the closed expression formulas in Therorem 1 illustrates a basic feature of the model — larger variance increases the discrepancy between *μ* and 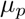. It is also robust to non-normality, as we will see in section 3.2. In case we have enough observations to motivate the *Z*-test, we perform a simple *Z*-test and obtain a *p*-value based on a two sided test (both deletions and insertions are tested for) using the *z*-statistic

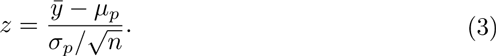

We refer to the null-hypothesis test using (3) as 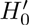. Thus, we have derived a different distribution under the null-hypothesis which we advocate should be used instead of *H*_0_. In case we have few observations (more often over insertions), approximation with the *Z*-test is poor. To get an exact test we would need to derive the distribution of 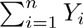, for *n* observations 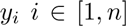. This could improve power to detect insertions, but we refrain from studying this in the present paper.

## 3 Results

We discuss why modeling bias contributes to making deletion calls more frequent than insertions calls in section 3.1. In section 3.2 we show that our corrected hypothesis agrees with biological data, and in section 3.3 that how indel detection is affected in CLEVER when our null-hypothesis is inserted.

### 3.1 Bias between detection of deletions and insertions

As donor fragments need to span over insertions 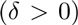, and this probability is 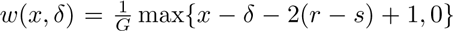 according to the OFL model, it is less likely that such fragments will be observed, as *δ* grows. We will therefore have a lower sample size over insertions in general. This naturally gives less power to detect an insertion compared to a deletion of similar size. However, methods using *H*_0_ have less power than necessary. Firstly, as 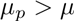, this gives too many significant upper quantile *p*-values (deletions) and too few significant lower quantile *p*-values (insertions). The difference in significance of observing a fragment of size 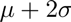 compared to observing a fragment of size 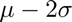 under *H*_0_, compared to the when observed under 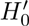 is seen in Figure 1b. Secondly, the positive skew of the OFL distribution (Figure 1b) makes a Z-test approximation less powerful compared to an exact test, especially for small sample sizes — as is more likely for insertions.

### 3.2 Evaluating the accuracy of 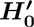

We evaluated the accuracy of our null-hypothesis on three mate pair libraries. We used a mate pair library from *Rhodobacter sphaeroides* from [21] denoted *rhodo*, a mate pair library from *Plasmodium falciparum* used in [13] denoted *plasm*, and mate-pair data from a human individual in the CEPH 1463 family-trio^10^. For the human dataset we aligned the reads to the complete human genome, but limited analysis to chromosome 13. We call this dataset *hs13*. Table 1 shows information about the datasets. Recall from section 2.4 that 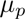 and 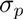 are the true mean and standard deviation of fragment lengths over a position that does not contain a variant. Let 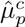 and 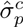 refer to the estimated quantity of 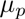 and 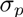 from the closed formulas in Theorem 1. Similarly, let 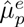 and 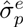 be the estimates of 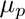 and 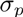 by using an empirical distribution of *f*(*x*) (estimated from a sample) and summing up the probabilities in equation (2) with 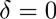. Estimates and observed values are shown in Table 1. It is our assumption that an overwhelming majority of positions are variant free^11^. Thus, we expect a model that fits data should give a uniformly distributed *p*-value distribution. Our observations are summarized below.

**Table 1:** Library information. Reads were aligned with BWA-MEM [16] version 0.7.12 with default parameters. Physical coverage is *c*, for all reads, and *c* (pp), for restricted proper pairs, i.e., read pairs that have both mates mapped in correct orientation and within a distance that depends on a statistical filtering of outliers based on the library distribution. The filtering bounds were roughly 10000, 6000, and 14000 bp for rhodo, plasm, and hs13 respectively. *μ* and *δ* are the mean and standard deviation of the full fragment length distribution. True mean insert-size and standard deviation over a position on the genome, 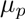 and 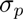 (calculated as the average over all positions in the genome) and predictions with closed formula, 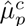 and 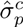 and exact calculation, 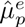 and 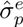.

| Organism | $r$ | $c$ | $c$ (pp) | $\mu$ | $\sigma$ | $\mu_p$ | $\sigma_p$ | $\hat{\mu}_p^e$ | $\hat{\sigma}_p^e$ | $\hat{\mu}_p^c$ | $\hat{\sigma}_p^c$ |
| --- | --- | --- | --- | --- | --- | --- | --- | --- | --- | --- | --- |
| rhodo | 101 | 43 | 34.5 | 2640 | 1390 | 3480 | 1534 | 3446 | 1526 | 3434 | 1143 |
| plasm | 75 | 4.9 | 4.2 | 2955 | 524 | 3056 | 511 | 3056 | 517 | 3053 | 515 |
| hs13 | 80 | 11.1 | 9.0 | 2947 | 1454 | 3688 | 1780 | 3719 | 1806 | 3705 | 1241 |

**Predicting 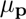**: 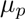 is estimated very well by both 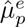 and 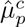 compare Figures 2d for hs13, and Figure 3b and 3d in Appendix for rhodo and plasm respectively with the estimated values in Table 1. Hence, testing 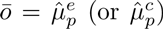 as in 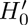 introduce symmetrical cumulative distribution function (CDF) values, Figure 2f, compared to CDF based on testing 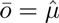 where all values are distributed around 1.0 — suggesting significant deletions, see Figure 2c.

**Fig 2:**
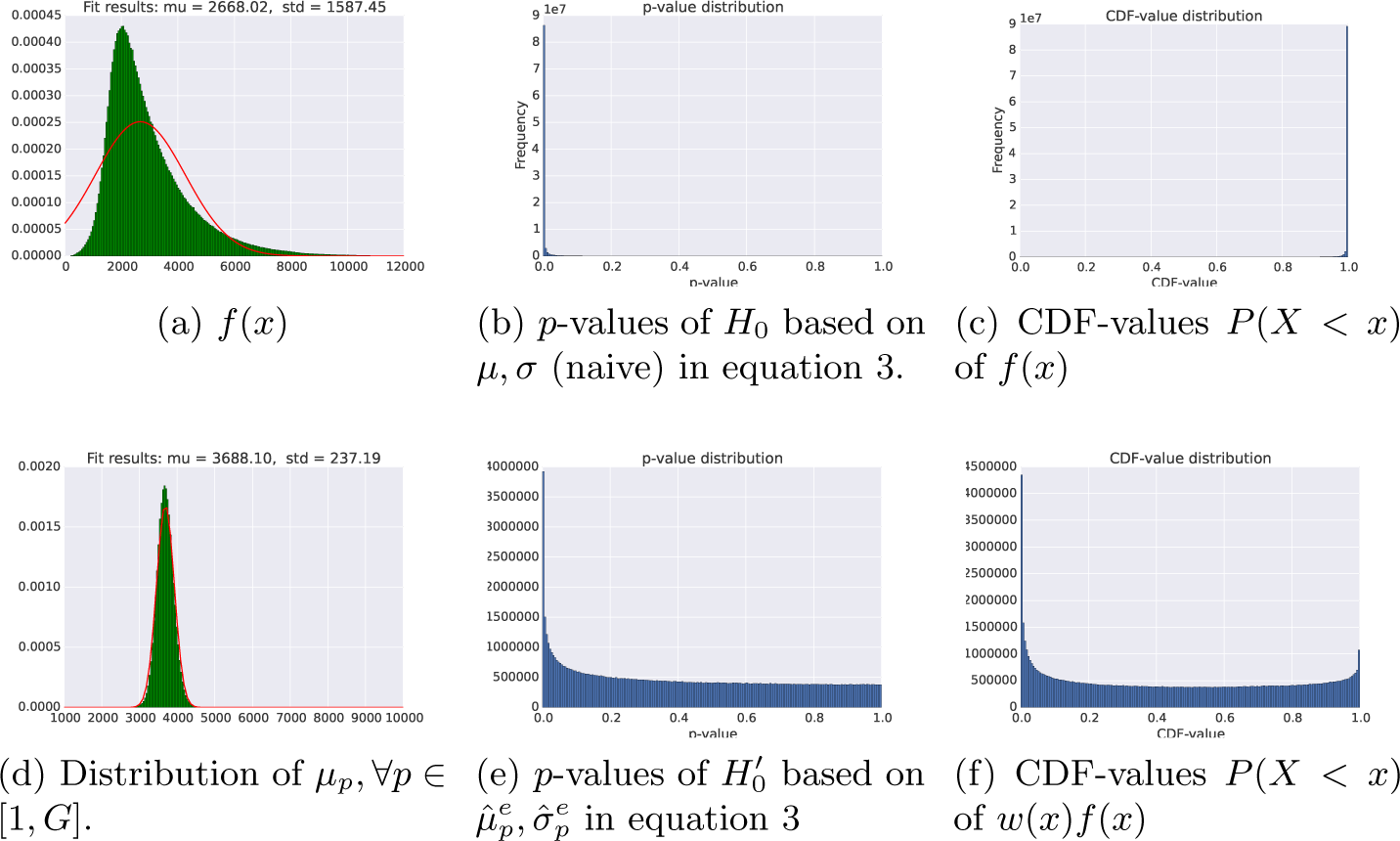
(a) The fragment length distribution *f*(*x*) for the hs13 dataset and the red line is a best fit of a truncated normal distribution. *f*(*x*) deviates significantly from a normal distribution. Although the mean of *f*(*x*) is 2947, the average observed fragment length over position 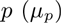 over all positions on hs13 shows that most values occur between 3000-4500 bp with the average around 3688 bp (d) — as approximately predicted from 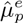 and 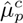. Figure (b) and (c) shows the *p*-value distribution and CDF values from using *H*_0_ (i.e., using *μ* and *σ*). Figure (e) and (f) shows the *p*-value distribution and CDF values from using 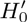 (i.e., using 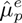 and 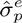).

**Fig. 3:**
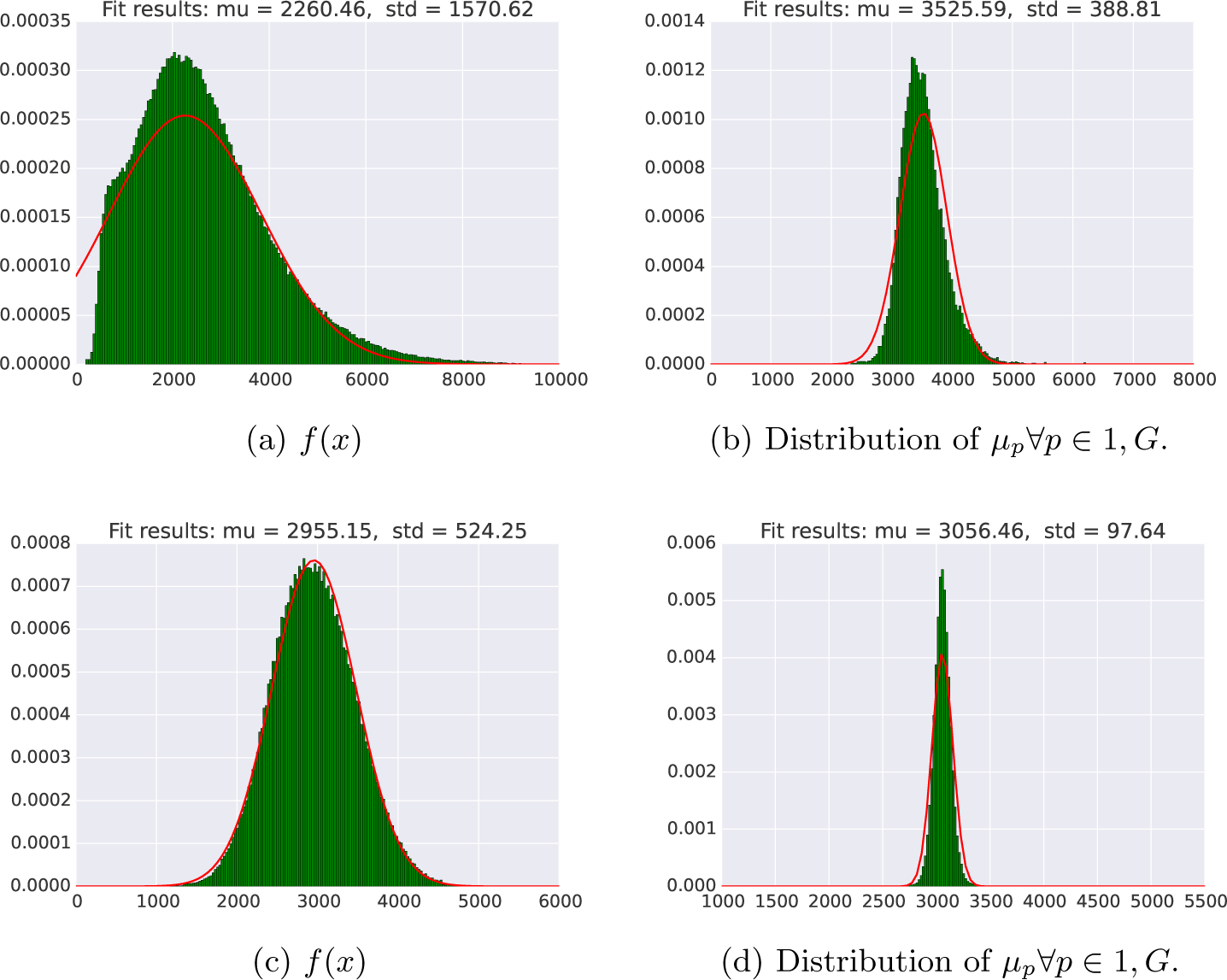
(a) rhodo library fragment size distribution. (b) Histogram over observed mean fragment length 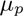 for all positions in the rhodo genome. (c) plasm library fragment size distribution. (d) Histogram over observed mean fragment length 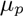 for all positions in the plasm genome.

**Predicting 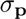**: The closed formula predictions of 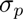 works best if *f*(*x*) is normal (plasm). For rhodo and hs13, 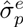 and 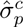 differs significantly and 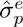 should be used, compare 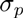 with 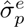 and 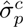 in Table 1.

***p*-values**: The *p*-value distribution (ideally uniform) greatly improves with 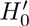 (Figure 2e) compared to *p*-values obtained with *H*_0_ (Figure 2b). Abnormalities in the *p*-values are most likely explained by: alignment artifacts (some regions are more difficult aligning to), fragment length bias [1,19], coverage bias from GC-content, and in some cases, real variants, see evaluation of hs13 dataset in section 5.7 of the Appendix. Similar *p*-value distributions are obtained on rhodo and plasm genome (data not shown) — that should not contain any variants — indicating that most of the enrichment of low *p*-values on hs13 is explained by any of the former three causes.

### 3.3 Implementing the corrected null-hypothesis in CLEVER

In this section we illustrate as a proof-of-concept how the corrected hypothesis 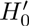 (with 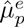 and 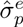) balances the ratio between detected insertions and deletions. We applied 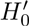 in CLEVER (v 1.1). *However, we want to emphasize that we did not tailor the statistical tests as needed to fit the assumptions made by their particular method. This limits the performance improvement*. To further improve results with CLEVER, we would need to (1) implement exact tests for few observations — giving more power to detect insertions, (2) use the OFL-model for CLEVER’s discovery of positions to study, (3) based on our model adjust CLEVER’s methods to handle, *e.g.*,heterozygous variants and controlling the false discovery rate. This would require additional modeling and significant restructuring of the code and we do not consider it here. Our aim here is only to illustrate how the simple adjustment of inserting 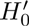 instead of *H*_0_ in CLEVER has a significant impact on the output. We investigated how the replacement of 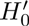 instead of *H*_0_ changed variant calls from CLEVER on hs13, rhodo and plasm as well with ideal condition simulated data denoted 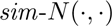 (full simulated results in Appendix section 5.6). For simulated variants, similarly to [17], a prediction is classified as a true positive (TP) if the breakpoint prediction is not further than one mean insert size (*i.e.*, at most 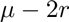) away from the true breakpoint. Otherwise it is classified as a false positive (FP). All variant calls on rhodo and plasm that are not from simulated variants are assumed to be false positives.

Because hs13 likely harbors true variants, we used annotated variants from dbVar [7], together with manual inspection in BamView [3], to assess if *hits* are true or false positives. For a deletion call in CLEVER with start and end coordinates *p*_s_, *p*_e_ and a deletion in dbVar with coordinates *q*_s_, *q*_e_, we let 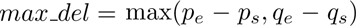 and 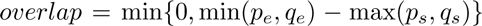. We let *hit_valwe = overlap/max_del and a call is a hit* if 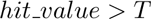, where 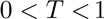 is a threshold. Because dbVar contains a large amount of annotated variants from several individuals and CLEVER produces many calls under *H*_0_, roughly 173, 106 and 40 hits are expected by chance with *T* = 0. 25, 0.5, 0. 75 (estimation in section 5.3), which is similar numbers to the observed hits from CLEVER: 226, 109 and 31 respectively under *T* = 0. 25, 0. 5, 0.75. We therefore further manually evaluated the hits produced with *T* = 0. 75 by looking for coverage drop and accumulation of softclips near each breakpoint. This gave us Estimated True Positives (ETP) as a rough measure of the TP rate for hs13. Therefore, we report ETP and Total Calls (TC) for hs13 in Table 2, contrary to simple TP and FP for the other data sets where we have the ground truth.

**Table 2:** Insertions and deletions called with CLEVER using *H*_0_ and 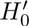. Column *δ* contains the size of 50 insertions and deletions, simulated on the reference genomes by either deleting or inserting a *δ* bp sequence on the reference. A “0” indicates that the original biological dataset was used.

| Dataset | $\delta$ | $H_0$ | | | | $H'_0$ | | | |
| --- | --- | --- | --- | --- | --- | --- | --- | --- | --- |
|  |  | TP (del/ins) |  | FP (del/ins) |  | TP (del/ins) |  | FP (del/ins) |  |
| plasm | 0 | 0 | (0/0) | **20** | (6/14) | 0 | (0/0) | 27 | (6/21) |
|  | 2000 | 89 | (50/39) | **22** | (8/22) | 89 | (50/39) | 38 | (8/30) |
| rhodo | 0 | 0 | (0/0) | 78 | (78/0) | 0 | (0/0) | **18** | (14/4) |
|  | 2000 | 49 | (49/0) | 54 | (54/0) | **57** | (45/12) | **13** | (9/4) |
| sim-N(500,75) | 75 | **94** | (94/0) | 0 | (0/0) | 78 | (62/16) | 0 | (0/0) |
|  | 100 | 110 | (100/10) | 0 | (0/0) | **188** | (100/88) | 0 | (0/0) |
|  |  | ETP (hits) |  | TC (del/ins) |  | ETP (hits) |  | TC (del/ins) |  |
| hs13 | 0 | **9** (31) |  | 1740 (1740/0) |  | 3 (4) |  | **8** (8/0) |  |

**Improvements**: From Table 2 and Figure 5 (Appendix) we see that CLEVER with *H*_0_ detects significantly more deletions than insertions of the same sizes. Using 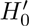, reduces this bias to some extent by increasing the detection of insertions across all data sets. CLEVER also returns significantly fewer false positive deletion calls with 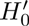, see rhodo Table 2 and 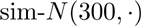. Even though *H*_0_ have more sensitivity in calling deletions on hs13, the signal disappears in the overwhelming amount of total calls, compare ETP and TC for *H*_0_ and 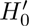 in Table 2.

**Deterioration**: A consequence of using 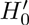 is fewer deletion calls, which unfortunately also removes some true positive deletion calls (see Figure 5 and Table 2). It also increases the FP insertion calls on plasm, see Table 2. We believe that calling variants with the plasm library carries additional difficulties due to its GC-poor genome sequence, such as positional fragment length bias [1,19].

Additional evidence that most calls with *H*_0_ on hs13 are FPs are found by comparing statistics on CLEVER’s deletion calls (Figure 6) and numbers reported in recent extensive studies [4,6]. For example, [4] provide frequency distributions for both previously discovered and new deletions on single genomes. Roughly 250 deletions have lengths over 1000 bp (inspection of plot). The simplifying assumption that large-indel distribution is uniform over chromosomes gives around 8 expected deletions^12^ in size range 1000 bp. This approximate number, and the fact that almost all calls were removed when using 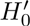 corroborates, that the vast majority (in the order of > 99%) of calls with *H*_0_ are FPs — likely a consequence of using 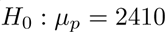 compared to the true value 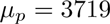.

## 4 Conclusions

We stated a probability distribution of observed fragment length over a position or region and derived a new null-hypothesis for detecting variants with fragment length, which is sound and agrees with biological data. Applied in CLEVER, our null-hypothesis detects more insertions and reduces false positive deletion calls. Results could be further improved by deriving an exact distribution instead of a *Z*-test and updating CLEVER's edge-creating conditions to agree with our model. The presented model, distribution, and null-hypothesis are general and could be used together with other information sources such as split reads, softclipped alignments, and read-depth information.

## Footnotes

5 With some modifications to account for heterozygous variants. Only reads that have enough overlap and similar fragment lengths are grouped together.

6 Under a normal distribution, 100 continuous observations are statistically equivalent to 158 binary observations for the best possible “cut point”, which is the mean. The loss of information becomes worse the further away the cut point is from the mean, e.g., 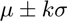, as *k* increases. In practice 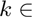 [3,6] in variant detection tools.

7 We call the side of the read that is closest to its mate “inner”.

8 A more informative prior could improve results, e.g., by fitting to the expected frequency and length of variants, studied in [4,6]. By tailoring the prior we could essentially obtain any specificity and sensitivity for a given indel size. We believe that is promising future work.

9 *n* to obtain sample mean 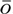, and log *t* to search the convex ML curve.

10 http://www.ebi.ac.uk/ena/data/view/ERR262996

11 Even small variants 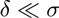 will not affect the model much.

12 Estimated as 250. 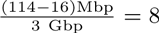, including compensation for the 16M N's at the start of the reference sequence for chr 13.

## 5 Appendix

### 5.1 Program versions and parameters

For CLEVER we used version v2.0-rc1 with parameters sorted, use_xa, -f, -w work.dir. For BWA we used BWA-mem version 0.7.12 with default parameters.

### 5.2 Rhodo and plasm libraries

### 5.3 Expected number of dbVar hits

Because there are many calls and annotated variants, we here provide a rough estimation of the number of expected hits at random. Let *k* be the number of variants in dbVar and *m* be the number of variant calls. For any pair of lengths of a deletion call and an annotated variant in dbVar under inspection, let *C*_min_ and *C*_max_ be the smallest and largest length in the pair, respectively. Let *C*_T_ be the total number of positions that two variants can be placed on, such that they will overlap with at least *TC*_max_ base pairs (an overlap of 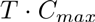 base pairs is required to be counted as a hit for threshold *T*, see section 3.3). Under the assumption that variant calls and annotated variants are randomly placed on the genome, we have that 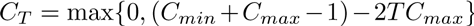. We now have that for *k* non-overlapping annotated variants of the *same size* and one variant call, the probability that a call will hit at least one annotated variant under threshold *T* is a simple summation of possible placements divided by genome size, *i.e.*, 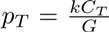. Furthermore, the expected number of hits (assuming calls are randomly generated from the genome) is obtained as 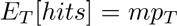. The limitation with this simplified formula is that it assumes fixed sizes of both calls and annotated variants when, in fact, they are random variables. We also assume that the variants are not overlapping.

Under these assumptions, we can now estimate the expected number of false hits based on numbers matching our data. For instance, we let a call be of size 2500 bp (most frequent size, see Figure 6), and we let the annotated variants all be 2000 bp (rough median estimation by inspecting variant lengths *v* with 250 < *v* < 8000, Figure 4). We chose 250 < *v* < 8000 because most variants outside this interval will get a low hit_valwe, as most calls are in this range Figure 6b. A rough estimation of the number of “non-redundant” (many annotated variants have identical start and end coordinates and some of them approximately the same start and stop) is 3000. We get this number by counting the number of variants that has 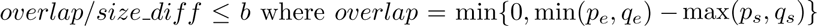 (from section 3.3) and 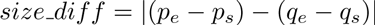. Thus, a low value suggests either a unique location or size compared any other variant, we get the *k* = 2747, 3217 and, 3504 for b = 1, 2, 3 respectively. Notice that these remaining variants are only used to get a rough estimation for *k* in this section. All variants are used to find *hits* as described in section 3.3. We get 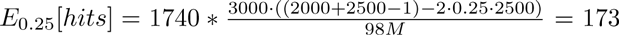 and similarly 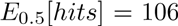, and 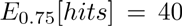. These are very rough estimations of the number of hits we could expect at random given *k* annotated variants and *m* variant calls which could be compared to the observed CLEVER hits 226, 109 and 31. Even though our calculation builds on many simplifications, the derived expected number of hits at random and the observed hits shows similar trends — suggesting that the majority of hits are expected at random.

**Fig. 4:**
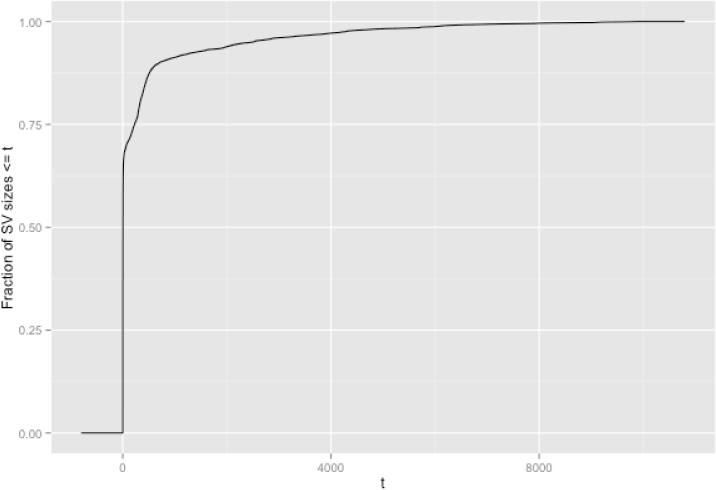
Cumulative length distribution of annotated variants in dbVar (>50 bp). Out of variants 250 <*v* <8000, 2000 seems to be the median length. The reason for looking at variants 250 < *v* < 8000 is that smaller or larger variants than this size will have very low hii_value as most calls are far from these cutoffs, Figure 6.

### 5.4 Reference implementation of the *p*-value evaluation

The implementation of this analysis is available from https://github.com/ksahlin/GetDistr/tree/develop/getdistr/assemblymodule. We want to emphasize that this is not intended to be a software for variant calling — but merely serves as a reference implementation.

To obtain the subset of reads over every position on which we calculate a *p*-value from we only need to read through a sorted bam-file twice (sampling from *f*(*x*) and calculating the metrics over each base pair). We use a window of positions on which read pairs it keeps in memory. Thus, the implementation for such an analysis has low time complexity and is scalable to a full human genome. The hs13 data takes around 1h and a maximum of 4Gb to process using one core (python code). This implementation can also be further extensively optimized.

### 5.5 Derivation of Result 1 and 2

From [22] we have 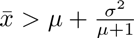 since 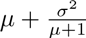 is the expected fragment length over any base pair. The greater sign comes from the lack of the constraint that at least *r* − *s* bases should be aligned on each side of *p*. Such constraint is needed in practice. For example, CLEVER uses *s* = 2 in its implementation which means that at least *r* − 2 base pairs from both reads must be located on respective sides of the variation. BreakDancer has no such criterion, but the criterion is then imposed on the read aligner being able to map at least *r* − *s* bases on respective sides. This gives the condition 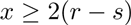.

Let 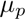 denote the mean of the distribution of reads spanning *p*. An exact value of 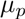 can be obtained for arbitrary distributions *f* by calculating the expected fragment length in equation 2 with 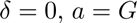 and 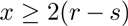. We can however give an accurate approximation of 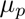 by letting 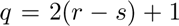 and substituting the 1's to q's in [22] (section 2.4, derivation of equation 2). We get Result 1 from this calculation. The derivation is identical, we therefore omit it here and only discuss why it's an accurate approximation.

The approximation is motivated as follows. The derivation in [22] (section 2.4) is assuming infinite support. Therefore, the above approximation is only accurate if the upper and lower boundaries are not located near high density regions of *f* (e.g. near the mode if 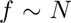). It is easy to motivate that *G* (the upper boundary) satisfies this. The lower boundary *q* is in practice also small enough to make the area between 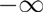 and *q* be negligible. The general conclusion that 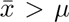 is already stated in [22]. Here, we also observe that 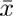 increases as the constraint 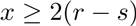 increases.

Similar to above, let *σ*_ρ_ be the standard deviation of the distribution of reads spanning a position *p*. Using the relation 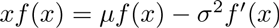, we have

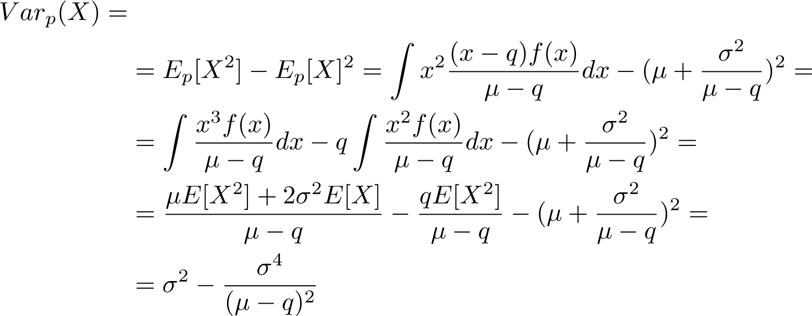

From this derivation. We immediately get the result in Theorem 1. The approximation of 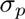 is following from the same assumptions as in the derivation of 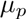 above and is a special case of the result in [23]. For hypotheses testing of variants with the assumptions above, 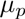 should be used in *H*_0_ and 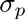 in the significance test.

### 5.6 CLEVER calls simulated data

We simulated 100 insertions and deletions respectively with sizes 10, 20, 30, 40, 50, 75, 100. We also simulated three different paired end libraries with *μ* = 300,400, 500 and 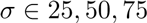 of 100bp error-free reads. All variations were on a distance of 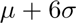 from each other and enough reads were generated so that CLEVER estimated *μ* and *σ* within 0.5 base pairs accuracy in all experiments — ideal conditions. Results are shown in Figure 5. We note that most of the variant sizes investigated here are too small to be detected with fragment length in cutoff based approaches without accepting a large amount of false positives. For example, the default cutoffs of considered fragment lengths in BreakDancer and Ulysses need to differ from *μ* with 3*σ* and 6*σ* respectively.

**Fig. 5:**
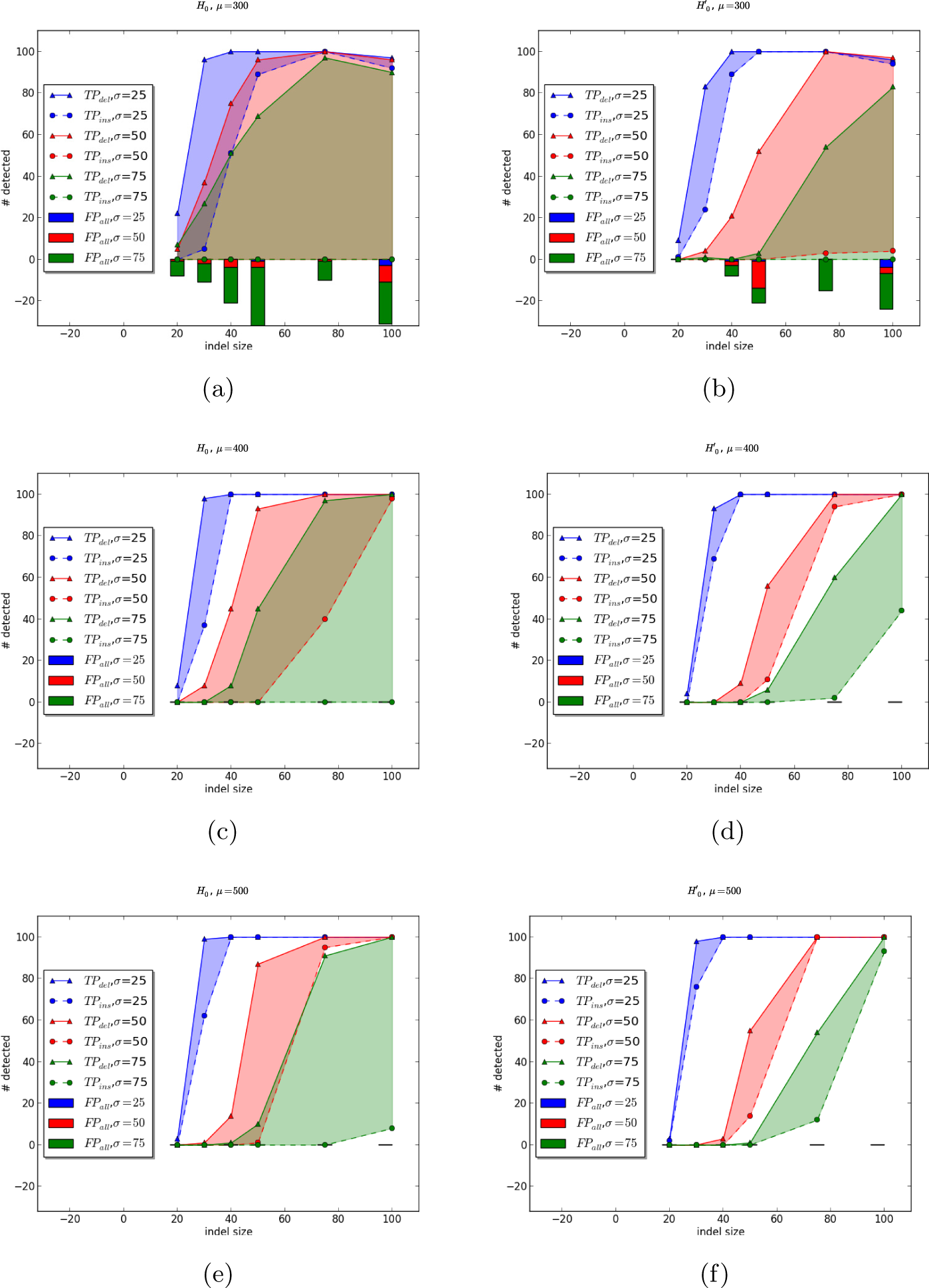
True and false positives for CLEVER when detecting insertions (dashed lines) and deletions (solid lines) of different sizes (*x*-axis) on a simulated genome with 100× (uniform) coverage from a normally distributed read pair library 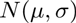. The insert size distribution was accurately inferred by CLEVER in all simulations. The colors indicate three different library widths, 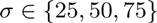. Shaded area displays difference between number of insertions and deletions detected, and in a well-balanced test this area should be small. The experiment is performed for 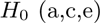 and 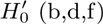.

### 5.7 CLEVER deletion length calls on hs13

**Fig. 6:**
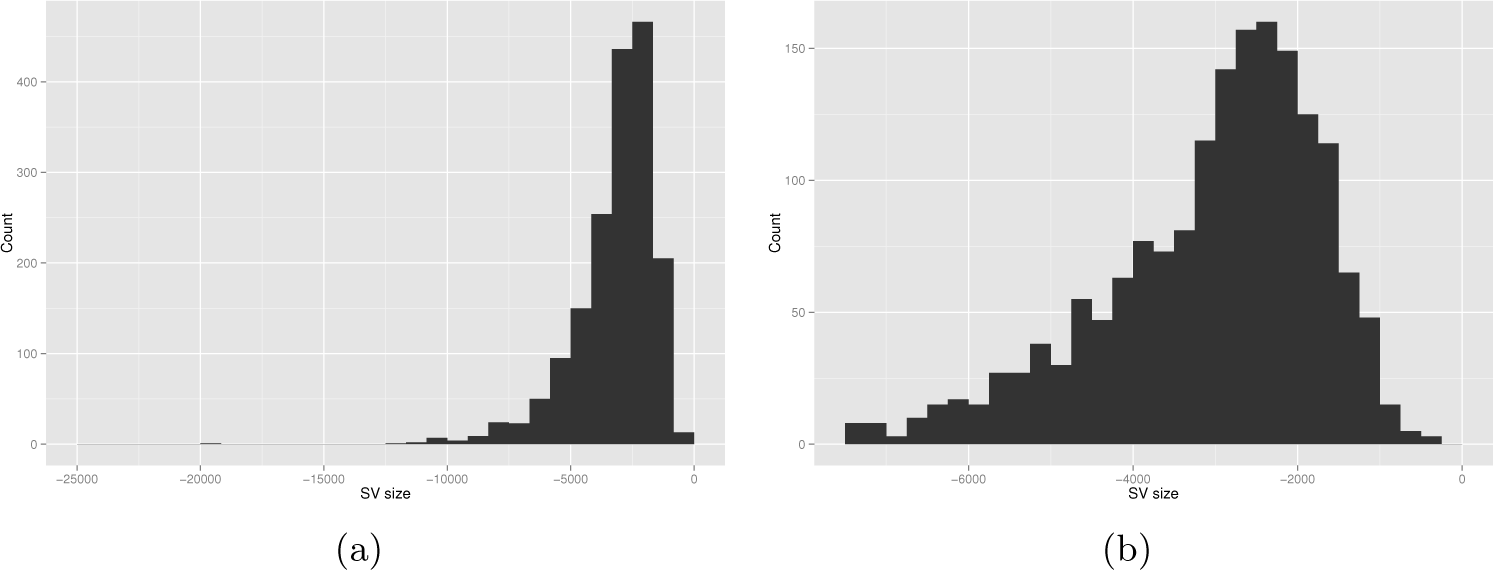
Length histogram of deletion calls on hs13. (a) Full histogram, and (b) a zoom-in of the region [-8000, 0].

